# ANME-2d anaerobic methanotrophic archaea differ from other ANME archaea in lipid composition and carbon source

**DOI:** 10.1101/558007

**Authors:** Julia M. Kurth, Nadine T. Smit, Stefanie Berger, Stefan Schouten, Mike S.M. Jettena, Cornelia U. Welte

**Author notes:** Correspondence: Julia Kurth, Department of Microbiology, Institute for Water and Wetland Research, Radboud University, Heyendaalseweg 135, 6525 AJ Nijmegen, The Netherlands;., Cornelia Welte,.

## Abstract

The anaerobic oxidation of methane (AOM) is a microbial process present in marine and freshwater environments. AOM is important for reducing the emission of the second most important greenhouse gas methane. In marine environments anaerobic methanotrophic archaea (ANME) are involved in sulfate-reducing AOM. In contrast, *Ca*. Methanoperedens of the ANME-2d cluster carries out nitrate AOM in freshwater ecosystems. Despite the importance of those organisms for AOM in non-marine environments not much is known about their lipid composition or carbon sources. To close this gap, we analyzed the lipid composition of ANME-2d archaea and found that they mainly synthesize archaeol and hydroxyarchaeol as well as different (hydroxy-) glycerol dialkyl glycerol tetraethers, albeit in much lower amounts. Abundant lipid headgroups were dihexose, monomethyl-phosphatidyl ethanolamine and phosphatidyl hexose. Moreover, a monopentose was detected as a lipid headgroup which is rare among microorganisms. Batch incubations with ^13^C labelled bicarbonate and methane showed that methane is the main carbon source of ANME-2d archaea varying from ANME-1 archaea which primarily assimilate dissolved inorganic carbon (DIC). ANME-2d archaea also assimilate DIC, but to a lower extent than methane. The lipid characterization and analysis of the carbon source of *Ca.* Methanoperedens facilitates distinction between ANME-2d and other ANMEs.

## Introduction

Methane is the second most important greenhouse gas on earth with an atmospheric methane budget of about 600 Tg per year (Conrad, 2009; Dean *et al.*, 2018). About 69% of methane emission into the atmosphere is caused by methanogenic archaea (Conrad, 2009). Fortunately aerobic and anaerobic methanotrophic microorganisms can oxidize methane back to carbon dioxide that is a 25-times less potent greenhouse gas than methane. The anaerobic oxidation of methane (AOM) is a microbial process present in marine and freshwater environments. AOM has first been described to be performed by a consortium of anaerobic methanotrophic archaea (ANME) and sulfate-reducing bacteria in microbial mats in the deep sea or in marine sediments (Hoehler *et al.*, 1994; Hinrichs *et al.*, 1999; Boetius *et al.*, 2000; Hinrichs and Boetius, 2002; Orphan *et al.*, 2002). ANME archaea are related to methanogens and oxidize methane by using the reverse methanogenesis pathway (Hallam *et al.*, 2004; Arshad *et al.*, 2015; McAnulty *et al.*, 2017; Timmers *et al.*, 2017). In addition to sulfate, also oxidized nitrogen compounds (Raghoebarsing *et al.*, 2006; Ettwig *et al.*, 2010; Haroon *et al.*, 2013) as well as iron and manganese (Beal *et al.*, 2009; Ettwig *et al.*, 2016; Cai *et al.*, 2018) can be used as electron acceptors within the AOM process.

Anaerobic methanotrophic archaea can be assigned to three distinct clusters within the Euryarchaeota, ANME-1, ANME-2 and ANME-3, which are related to the orders *Methanosarcinales* and *Methanomicrobiales* (Knittel and Boetius, 2009). The phylogenetic distance between the groups is quite large (16S rRNA gene sequence identity between 75-92%) (Knittel and Boetius, 2009). Most analysed members of the three ANME clades have been described to perform sulfate driven AOM in marine environments (Pancost *et al.*, 2001; Blumenberg *et al.*, 2004; Niemann and Elvert, 2008; Rossel *et al.*, 2008; Wegener *et al.*, 2008; Kellermann *et al.*, 2012). However, members of the ANME-2d cluster have not been found in consortia with sulfate reducers. Instead, ANME-2d archaea are the main players in nitrate-dependent AOM. Microorganisms conducting nitrate AOM have been enriched from anoxic freshwater sediments, digester sludge and rice paddies (Raghoebarsing *et al.*, 2006; Hu *et al.*, 2009; Arshad *et al.*, 2015; Vaksmaa, Guerrero-Cruz, *et al.*, 2017). Denitrifying AOM can either be conducted by a consortium of nitrate-reducing ANMEs, *Ca*. Methanoperedens sp., and nitrite reducing NC10 bacteria, *Ca.* Methylomirabilis sp. (Raghoebarsing *et al.*, 2006; Haroon *et al.*, 2013; Arshad *et al.*, 2015) or by a consortium of those ANME archaea and anammox bacteria (Haroon *et al.*, 2013). In those consortia *Ca.* Methylomirabilis sp. or anammox bacteria are important to reduce the toxic nitrite produced during nitrate AOM by *Ca*. Methanoperedens sp.

To understand the prevalence of anaerobic methane oxidation in past and present environments and identify the key players at different environmental sites, it is necessary to identify biomarkers for those organisms. As core lipids are much more stable than DNA over time, lipid biomarkers are a useful tool to trace microorganisms and therefore specific microbial processes back in time. Moreover, intact polar lipids are crucial to examine present microbial communities and to distinguish between different microorganisms (Ruetters *et al.*, 2002; Sturt *et al.*, 2004). Quite some information is available on core and intact polar lipids as well as on carbon assimilation in marine AOM consortia of ANME archaea and sulfate-reducing bacteria (Pancost *et al.*, 2001; Blumenberg *et al.*, 2004; Niemann and Elvert, 2008; Rossel *et al.*, 2008; Wegener *et al.*, 2008; Kellermann *et al.*, 2012). In contrast, lipids from one of the main players in denitrifying AOM, *Ca*. Methanoperedens sp., have hardly been studied: a preliminary study on the lipids of a culture containing *Ca*. Methanoperedens sp. and *Ca*. Methylomirabilis oxyfera only detected sn2-hydroxyarchaeol as the dominant lipid of the archaeal partner (Raghoebarsing *et al.*, 2006).

Besides the characterization of lipids in ANME archaea it is also pivotal to understand which carbon source those organisms use for biomass production. The main carbon assimilation pathway in methanogenic Euryarchaeota is the reductive acetyl-CoA pathway (Whitman, 1994; Berg et al., 2010). In this pathway a carbonyl group and a methyl group are combined to form acetyl-CoA. In archaea, acetyl-CoA is used for formation of membrane lipids via the isoprenoid compound geranylgeranylphosphate in the mevalonate pathway, although not all of the enzymes involved in this pathway are known with certainty (Koga and Morii, 2007; Matsumi *et al.*, 2011). An ether bond is formed between the glycerol-1-phosphate backbone and the isoprenoid side chains. Subsequently cytidine-diphosphate is attached and finally the unsaturated isoprenoid side chains are reduced to form diphytanylglycerol diether, also known as archaeol (Matsumi *et al.*, 2011).

The isotopic composition of lipids provides information on the carbon source used by the microorganism. The lipids of ANMEs involved in AOM are usually strongly depleted in ^13^C, with δ^13^C values ranging from –70 to –130‰ (Elvert *et al.*, 1999; Pancost *et al.*, 2000; Niemann and Elvert, 2008). Such low δ^13^C values of lipids have been explained by the assimilation of ^13^C-depleted methane carbon during methane uptake into biomass (Elvert *et al.*, 1999; Hinrichs *et al.*, 1999; Pancost *et al.*, 2000; Orphan *et al.*, 2002). Mixed assimilation of CH_4_ and CO_2_ has been reported for marine ANME-1, -2a, and -2b strains indicating that at least some ANME strains can use methane-derived carbon for biomass production (Wegener *et al.*, 2008). However, for ANME-1 it has been shown that methane oxidation is decoupled from the assimilatory system and that CO_2_-dependent autotrophy is the predominant mode of carbon fixation (Kellermann *et al.*, 2012). In general, ANME archaea seem to be able to assimilate both, methane and dissolved inorganic carbon, and the preferred carbon source for assimilation might vary between the different ANME clusters.

In this study, we performed analysis of the lipids from ANME-2d archaea and compared these with previous studies about different ANME lipids. Moreover, we analysed the incorporation of ^13^C-labelled methane and bicarbonate in lipids of these archaea to establish the carbon sources used for assimilation.

## Methods

### ANME-2d bioreactor operation and sampling for lipid analysis

For lipid analysis of *Ca*. Methanoperedens sp. two different bioreactors were sampled. One bioreactor contained archaea belonging to the ANME-2d clade enriched from the Ooijpolder (NL) (Arshad *et al.*, 2015; Berger *et al.*, 2017) and the other reactor ANME-2d archaea enriched from an Italian paddy field (Vaksmaa, Jetten, *et al.*, 2017). The anaerobic enrichment culture dominated by *Ca*. Methanoperedens sp. strain BLZ2 originating from the Ooijpolder (Berger *et al.*, 2017) was maintained in an anaerobic 10 L sequencing batch reactor (30°C, pH 7.3 ± 0.1, stirred at 180 rpm). The mineral medium consisted of 0.16 g/L MgSO_4_, 0.24 g/L CaCl_2_ and 0.5 g/L KH_2_PO_4_. Trace elements and vitamins were supplied using stock solutions. 1000 x trace element stock solution: 1.35 g/L FeCl_2_ x 4 H_2_O, 0.1 g/L MnCl_2_ x 4 H_2_O, 0.024 g/L CoCl_2_ x 6 H_2_O, 0.1 g/L CaCl_2_ x 2 H_2_O, 0.1 g/L ZnCl_2_, 0.025 g/L CuCl_2_ x 2 H_2_O, 0.01 g/L H_3_BO_3_, 0.024 g/L Na_2_MoO_4_ x 2 H_2_O, 0.22 g/L NiCl_2_ x 6 H_2_O, 0.017 g/L Na_2_SeO_3_, 0.004 g/L Na_2_WO_4_ x 2 H_2_O, 12.8 g/L nitrilotriacetic acid; 1000 x vitamin stock solution: 20 mg/L biotin, 20 mg/L folic acid, 100 mg/L pyridoxine-HCl, 50 mg/L thiamin-HCl x 2 H_2_O, 50 mg/L riboflavin, 50 mg/L nicotinic acid, 50 mg/L D-Ca-pantothenate, 2 mg/L vitamin B12, 50 mg/L p-aminobenzoic acid, 50 mg/L lipoic acid. The medium supply was continuously sparged with Ar:CO_2_ in a 95:5 ratio. Per day 30 mmol nitrate added to the medium were supplied to the bioreactor and were completely consumed. Methane was added by continuously sparging the reactor content with CH_4_:CO_2_ in a 95:5 ratio at a rate of 15 mL/min. The reactor was run with a medium turnover of 1.25 L per 12 h. A 5 min settling phase for retention of biomass preceded the removal of supernatant. Under these conditions nitrite was not detectable with a colorimetric test with a lower detection limit of 2 mg/L (MQuant test stripes, Merck, Darmstadt, Germany). Growth conditions and operation of the bioreactor containing ANME-2d archaea enriched from an Italian paddy field soil are described by Vaksmaa *et al*., 2017 (Vaksmaa, Jetten, *et al.*, 2017). Sampled material from both reactors was centrifuged (10000 x *g*, 20 min, 4°C) and pellets were kept at −80°C until subsequent freeze-drying and following lipid and isotope analysis.

### Analysis of the microbial community

For the Ooijpolder enrichment we performed whole genome metagenome sequencing. DNA extraction, library preparation and metagenome sequencing were performed as described before by Berger and co-workers (Berger *et al.*, 2017). Quality-trimming, sequencing adapter removal and contaminant filtering of Illumina paired-end sequencing reads were performed using BBTools BBDuk 37.76 (BBMap - Bushnell B. - sourceforge.net/projects/bbmap/). Processed paired-end reads were assigned to a taxon using Kaiju 1.6.2 (Menzel *et al.*, 2016) employing the NCBI BLAST non-redundant protein database (NCBI Resource Coordinators, 2016).

### Batch cultivation of ANME-2d bioreactor cell material for ^13^C-labelling experiment

60 ml bioreactor material of a *Ca*. Methanoperedens sp. BLZ2 culture enriched from the Ooijpolder (Arshad *et al.*, 2015; Berger *et al.*, 2017) were transferred with a syringe to a 120-ml serum bottle that had been made anoxic by flushing the closed bottle with argon gas for 10 min. Afterwards, the culture was purged with 90% argon and 10% CO_2_ for 5 min. 2.5 mM NaHCO_3_ and 18 ml methane (Air Liquide, Eindhoven, The Netherlands) were added. Except for the negative controls, either ^13^C-labelled methane (99 atom%; Isotec Inc., Matheson Trigas Products Division) or ^13^C-labelled bicarbonate (Cambridge Isotope Laboratories Inc., Tewksbury, USA) was used in the batch incubations. The bottles were incubated horizontally on a shaker at 30°C and 250 rpm for one or three days. All bottles contained sodium nitrate (0.6 mM) at the start of incubation and additional nitrate was added when the concentration in the bottles was close to 0, as estimated by MQuant (Merck, Darmstadt, Germany) test strips. The methane concentration in the headspace was measured twice a day by gas chromatography with a gas chromatograph (Hewlett Packard 5890a, Agilent Technologies, Santa Clara CA, US) equipped with a Porapaq Q 100/120 mesh and a thermal conductivity detector (TCD) using N_2_ as carrier gas. Each measurement was performed by injection of 50 μl headspace gas with a gas-tight syringe. With this technique a decrease in methane concentration from ∼24 % to ∼20 % within three days of incubation was observed. After batch incubation, cell material was centrifuged (10000 x *g*, 20 min, 4°C) and pellets were kept at −80°C until subsequent freeze-drying and following lipid and isotope analysis. It has to be considered that cultures with ^13^C-labelled bicarbonate also contained ^12^C derived from CO_2_. Having in mind that 10% of CO_2_ were added to the culture (pCO_2_= 0.1 atm) the CO_2_ concentration in the solution was calculated to be about 3.36 mM by use of the equation [CO2]_aq_= pCO_2_/k_h_ (Henry constant (k_h_) is 29.76 atm/(mol/L) at 25°C). Therefore, it has to be assumed that about half of the carbon in the cultures where ^13^C labelled bicarbonate was added derived from ^12^C-CO_2_ dissolved in the medium after gassing with a mixture of 10% CO_2_/90% Ar gas.

### Lipid extraction and analysis

#### Bligh and Dyer extraction

Lipids of freeze-dried biomass (between 20 and 70 mg) were extracted by a modified Bligh and Dyer method as described by Bale et al. (Bale *et al.*, 2013) using a mixture of methanol, dichloromethane and phosphate buffer at pH 7.4 (2:1:0.8 v/v/v). After ultrasonic extraction (10 min) and centrifugation the solvent layer was collected. The residue was re-extracted twice. The combined solvent layers were separated by adding additional DCM and phosphate buffer to achieve a ratio of MeOH, DCM and phosphate buffer (1:1:0.9 v/v/v). The separated organic DCM layer on the bottom was removed and collected while the aqueous layer was washed two more times with DCM. The combined DCM layer was evaporated under a continuous stream of nitrogen.

#### Acid hydrolysis

Head groups of archaeal lipids were removed using acid hydrolysis. About 20 mg freeze-dried biomass was hydrolyzed with 2 ml of a 1.5 N HCl/MeOH solution and samples stirred for 2 h while heated at 130°C with a reflux system. After cooling, the pH was adjusted to pH 4-5 by adding 2 N KOH/MeOH solution. 2 ml DCM and 2 ml distilled H_2_O were added. The DCM bottom layer was transferred to a new vial and the MeOH/H_2_O layer washed twice with DCM. Combined DCM layers were dried over a Na_2_SO_4_ column and the solvent removed by evaporation under a stream of nitrogen.

#### BF3 methylation and silylation

For gas chromatography (GC) analysis aliquots of the acid hydrolyzed samples were methylated using 0.5 ml of BF_3_-methanol and react for 10 min at 60°C in a oven. 0.5 ml H_2_O and 0.5 ml DCM were added to the heated mixture to separate the DCM and aqueous layers. Samples were mixed, centrifuged and the DCM-layer taken off and collected. The water layer was washed three more times with DCM. The combined DCM-layer were evaporated under a N_2_ stream and water was removed by use of a MgSO_4_ column. After dissolving the sample in ethyl acetate, the extract was cleaned over a small silicagel column and lipids were eluted with ethyl acetate. The extract was dried under N_2_. For GC analysis, extracts (0.3 to 0.5 mg) were dissolved in 10 μl pyridine and 10 μl BSTFA. Samples were heated for 30 min at 60°C and afterwards diluted with ethyl acetate to 1 mg/ml.

#### GC-MS

Gas chromatography linked to mass spectrometry (GC-MS) was performed with a 7890B gas chromatography system (Agilent) connected to a 7000 GC/MS Triple Quad (Agilent). The gas chromatograph was equipped with a fused silica capillary column (25 m x 0.32 mm) coated with CP Sil-5 CB (0.12 μm film thickness) and a Flame Ionization Detector (FID). Helium was used as the carrier gas. The samples were injected manually at 70°C via an on-column injector. The oven temperature was programmed to a temperature increase from 70 to 130°C with 20°C/min and a further increase to 320°C with 4°C/min to, 320°C was held for 10 min. The mass range of the mass spectrometer was set to scan from m/z 50 to m/z 850.

#### GC-IRMS

Gas chromatography coupled to isotope-ratio mass spectrometry (GC-IRMS) was performed on a TRACE 1310 Gas Chromatograph (Thermo Fisher Scientific) interfaced with a Scientific GC IsoLink II Conversion Unit connected to an IRMS DELTA V Advantage Isotope-ratio mass spectrometer (Thermo Fisher Scientific). The gas chromatograph was equipped with a fused silica capillary column (25 m x 0.32 mm) coated with CP Sil-5 CB (0.12 μm film thickness). Helium was used as the carrier gas. The samples were injected at 70°C via an on-column injector. The oven temperature was programmed to a temperature increase from 70 to 130°C with 20°C/min and a further increase to 320°C with 4°C/min, 320°C was held for 10 min. δ^13^C values were corrected for methyl group derived from BF_3_ methanol in case of carboxylic acid group (bacterial lipids) and methyl groups derived from BSTFA in case of hydroxyl groups (mainly archaeal lipids). Averaged δ13C values are based on experimental triplicates, but not on analytical duplicates.

#### UHPLC-APCI-TOF-MS

About 0.4 to 0.8 mg of the acid hydrolyzed lipid extract was dissolved in a mixture of hexane/isopropanol 99:1. Extracts were filtered by use of a 0.45 μm, 4 mm diameter PTFE filter. About 2 mg per ml core lipid containing extracts were used for analysis by ultra-high performance liquid chromatography linked to time-of-flight atmospheric pressure chemical ionization mass spectrometry using a (UHPLC-APCI-TOFMS). Core lipid analysis was performed on an Agilent 1260 Infinity II UHPLC coupled to an Agilent 6230 TOF-MS. Separation was achieved on two UHPLC silica columns (BEH HILIC columns, 2.1 x 150 mm, 1.7 μm; Waters) in series maintained at 25°C. The injection volume was 10 μl. Lipids were eluted isocratically for 10 min with 10% B, followed by a linear gradient to 18% B in 15 min, then a linear gradient to 30% B in 25 min, then a linear gradient to 100% B in 30 min, and finally 100% B for 20 min, where A is hexane and B is hexane: isopropanol (9:1). Flow rate was 0.2 ml/min and pressure 400 bar. Total run time was 120 min with a 20 min re-equilibration. Settings of the ion source (APCI) were as followed: gas temperature 200°C, vaporizer 400°C, drying gas 6 l/min, nebulizer 60 psig. The lipids were identified using a positive ion mode (600–1400 m/z). 4mm PTFE filter (polypropylene…).

#### UHPLC-ESI-MS

0.3 to 0.7 mg of Bligh and Dyer sample was dissolved in an injection solvent composed of hexane/isopropanol/water (72:27:1;v/v/v) and filtered through a 0.45 μm regenerated cellulose filter with 4 mm diameter prior to analysis by ultra-high performance liquid chromatography linked to ion trap mass spectrometry using electrospray ionization (UHPLC-ESI-MS). UHPLC separation was conducted on an Agilent 1200 series UHPLC equipped with a YMC-Pack Diol-120-NP column (250 x 2.1 mm, 5 μm particle size) and a thermostated autoinjector, coupled to a Thermo LTQ XL linear ion trap with Ion Max source with electrospray ionization (ESI) probe (Thermo Scientific, Waltham, MA). Solvent A contained 79% hexane, 20% isopropanol, 0.12% formic acid, 0.04% ammonium and solvent B 88% isopropanol, 10 % H_2_O, 0.12% formic acid, 0.04% ammonium. Lipids were eluted with 0% B for 1 min, a linear gradient from 0 to 34% B in 17 min, 34% B for 12 min, followed by a linear gradient to 65% B in 15 min, 65% B for 15 min and finally a linear gradient to 100% B in 15 min. The IPLs were identified using a positive ion mode (m/z 400–2000) and a collision energy of 35 eV.

## Results and Discussion

### Analysis of microbial community

We performed phylogenetic analysis of the microbial community in the ANME-2d enrichment originating from the Ooijpolder. Twenty-three percent of the reads were assigned to *Ca*. Methanoperedens sp. strain BLZ2, 33% to *Ca*. Methylomirabilis sp. acting as nitrite scavenger, 8% to Alphaproteobacteria, 6% to Gammaproteobacteria, 5% to Betaproteobacteria 1% to Deltaproteobacteria, 3% to Terrabacteria, 3% to Sphingobacteria and 1% to Planctobacteria. The only archaeon in the bioreactor was *Ca*. Methanoperedens sp. strain BLZ2. Analysis of the microbial community in the Italian paddy field ANME-2d enrichment has been described by Vaksmaa et al., 2017. For this bioreactor a similar proportion (22%) of ANME-2d archaea, in this case *Ca*. Methanoperedens sp. strain Vercelli, was detected. In this study we mainly show the results derived from lipid analysis of the *Ca*. Methanoperedens sp. BLZ2 enrichment originating from the Ooijpolder (Arshad *et al.*, 2015; Berger *et al.*, 2017). However, the results deriving from a *Ca*. Methanoperedens sp. Vercelli enrichment originating from Italian paddy field soil (Vaksmaa, Jetten, *et al.*, 2017) look very similar, indicating that our results are not dependent on the strain or the environment from which the strain was enriched.

### Core lipids of *Ca*. Methanoperedens sp

To analyze the lipids of ANME-2d archaea, biomass from a bioreactor containing *Ca*. Methanoperedens sp. BLZ2 enrichment was sampled and core lipid analysis with GC-MS and UHPLC-APCI-TOF-MS was performed.

GC analysis of the core lipids released by acid hydrolysis showed that of the microbial community harbored bacterial fatty acids and isoprenoidal archaeal lipids (Figure 1). We detected the typical membrane lipids of *Ca*. Methylomirabilis sp., namely 10-methylhexadecanoic acid (10MeC_16:0_) and its monounsaturated variant (10MeC_16:1Δ7_) (Kool *et al.*, 2012). The archaeal isoprenoids were predominantly composed of archaeol with lower amounts of sn2-hydroxyarchaeol and sn3-hydroxyarchaeol as well as two monounsaturated archaeols (Nichols and Franzmann, 1992). Monounsaturated archaeol has already been described to be present in samples of archaea associated with anaerobic methane oxidation in marine environments (Pancost *et al.*, 2001; Blumenberg *et al.*, 2005). However, the monounsaturated archaeol might be produced from hydroxyarchaeol during acidic treatment of the lipids and therefore might not be part of native membrane lipid structures (Ekiel and Sprott, 1992). Alternation of the hydroxyarchaeol structure caused by different reaction conditions during lipid treatment has also been shown by Hinrichs and co-workers (Hinrichs *et al.*, 2000). On the other hand, monounsaturated archaeols have also been described for *Halorubrum lacusprofundi* (Franzmann *et al.*, 1988; Gibson *et al.*, 2005), *Methanopyrus kandleri* (Nishihara *et al.*, 2002), *Methanococcoides burtonii* (Nichols and Franzmann, 1992; Nichols *et al.*, 1993), even if using mild alkaline hydrolysis instead of acidic treatment for lipid extraction (Nishihara *et al.*, 2002).

**Figure 1:**
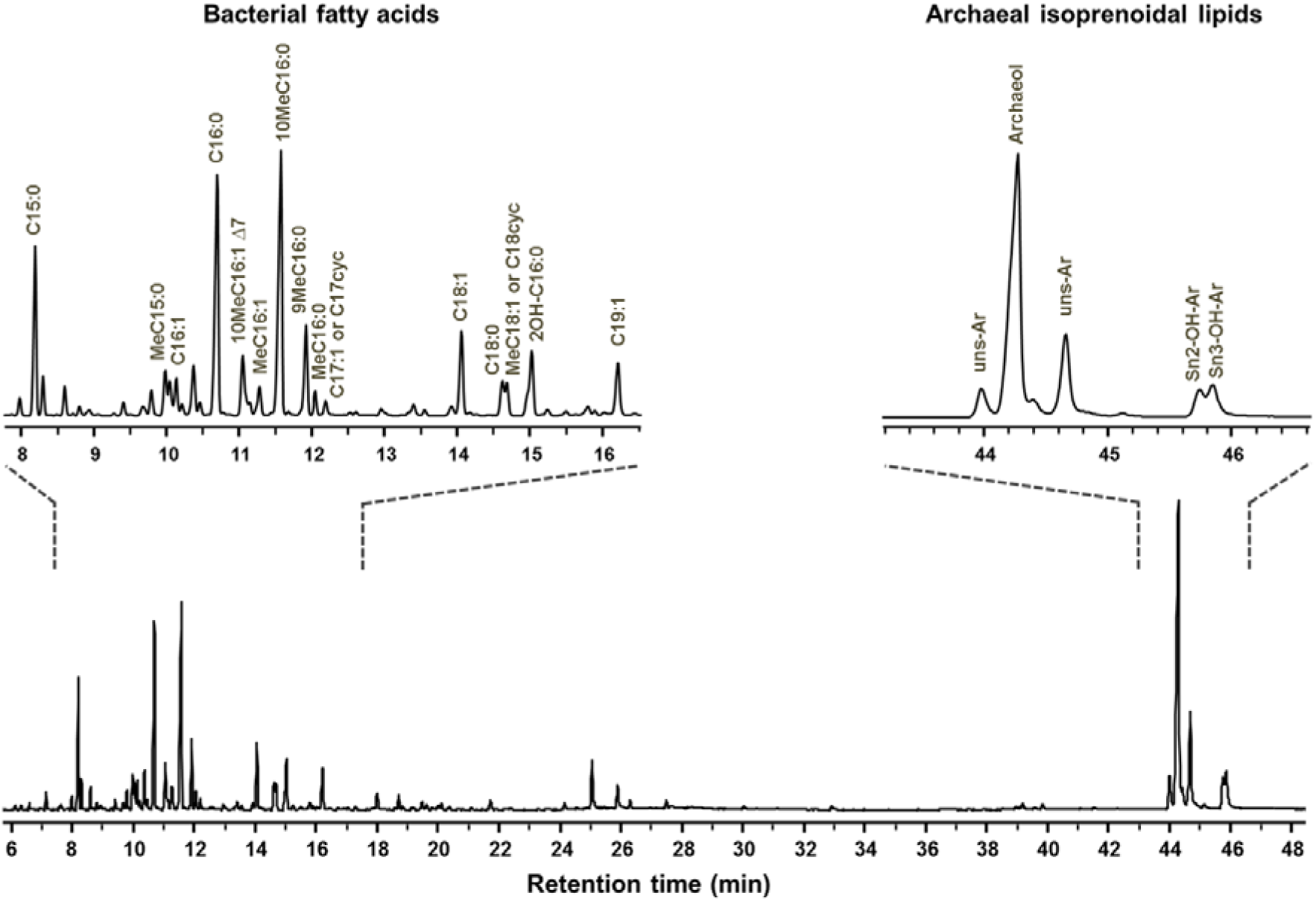
Gas chromatogram of core lipids released by acid hydrolysis from *Ca*. Methanoperedens sp. (ANME-2d) enrichment. The enriched biomass of ANME-2d originates from the Ooijpolder (NL) (Arshad et al., 2015). Enlarged inserts show the TIC (total ion chromatogram) of the bacterial and archaeal lipids. The most abundant compounds are annotated with their compound name and following abbreviations: Uns-Ar = monounsaturated archaeol, OH-Ar = hydroxyarchaeol.

One possibility to distinguish between the different ANME groups is the sn2-hydroxyarchaeol to archaeol proportion (Blumenberg *et al.*, 2004). For ANME-1 this ratio is described to be 0-0.8, for marine ANME-2 1.1 to 5.5 and for ANME-3 within the range of ANME-2 (Blumenberg *et al.*, 2004; Niemann *et al.*, 2006; Nauhaus *et al.*, 2007; Niemann and Elvert, 2008). In our study with the non-marine ANME-2d archaea we observed a sn2-hydroxyarchaeol to archaeol ratio of around 0.2. As mentioned before, the monounsaturated archaeol species might be an artefact of hydroxyarchaeol. If the monounsaturated archaeols are added to that of the sn2-hydroxyarchaeol abundance, the ratio would still be only around 0.3. That means that the hydroxyarchaeol to archaeol ratio of ANME-2d is more similar to that of ANME-1 archaea than to that of other ANME-2 or ANME-3 archaea. Members of the related methanogen order *Methanomicrobiales* only contain archaeol and GDGT-0 in their membranes but not hydroxyarchaeol (Koga *et al.*, 1998). In the order *Methanosarcinales* the lipid composition varies between the different members. Most strains produce archaeol and hydroxyarchaeol, but the ratio differs and also the type of hydroxyarchaeol isomer varies. *Methanosarcinaceae* mainly produce the sn2-isomer, whereas *Methanosaetaceae* mainly produce the rare sn3-isomer (Koga *et al.*, 1998).

Subsequently, UHPLC-APCI-TOF-MS analysis of the lipid extract was conducted in order to obtain information about the tetraether lipids. This revealed that the relative abundance of archaeol was two times higher than that of glycerol dialkyl glycerol tetraethers (GDGTs) (Table 1). Moreover, several types of GDGTs were present in the enrichment. GDGTs contained either no (GDGT-0), one (GDGT-1) or two (GDGT-2) cyclopentane rings and about 64% of the GDGTs were hydroxylated (OH-GDGTs). The most abundant GDGTs were GDGT-0 with 6% and di-OH-GDGT-2 with 5% of total lipids. In conclusion, ANME-2d archaea synthesize various core-GDGTs, however archaeol and its homologues are the main isoprenoidal core-lipids in this enrichment.

**Table 1:**
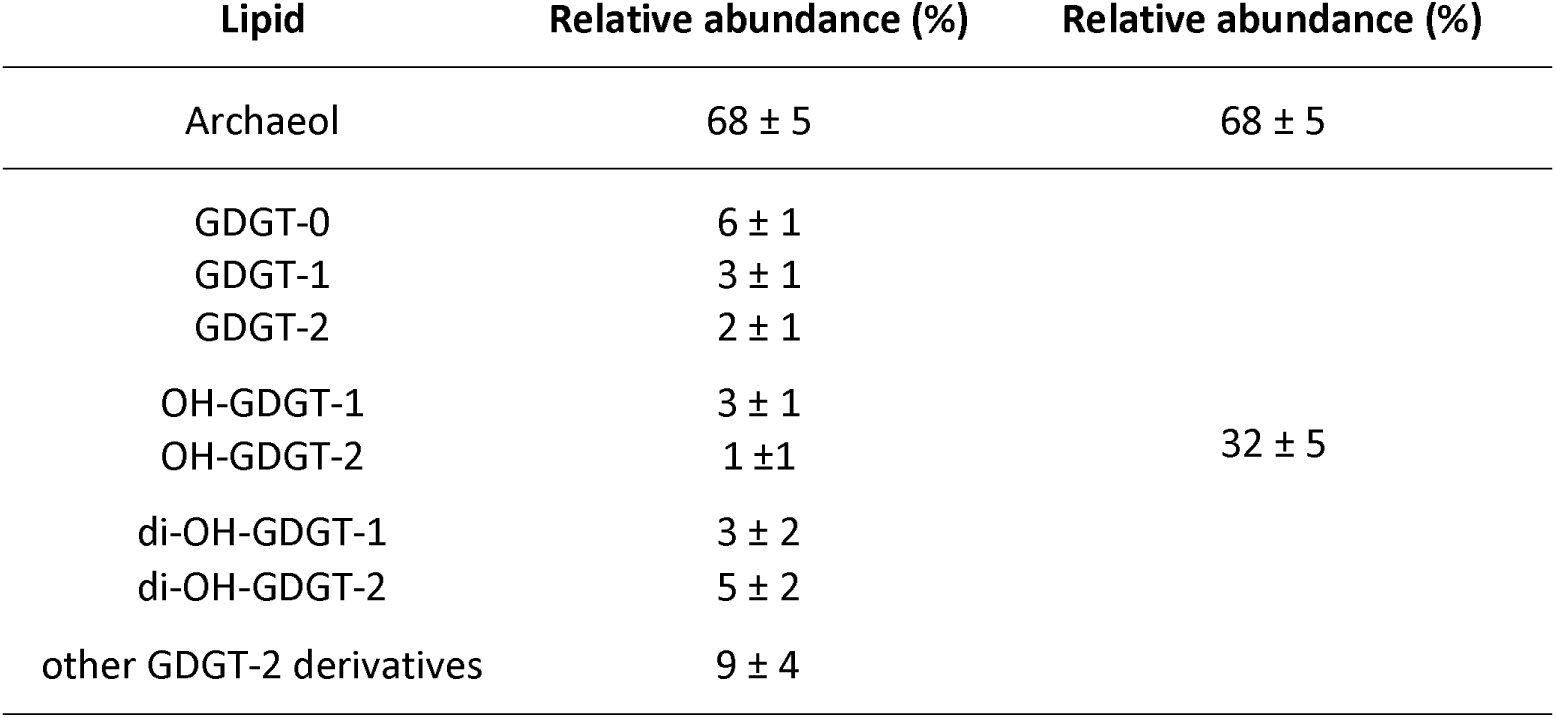
Abundance of archaeol and GDGTs of *Ca*. Methanoperedens sp. Lipid extraction was performed in quadruplicates, error is given as standard deviation. For calculation of the relative abundance of archaeol also peaks derived from archaeol artefacts created during the experimental procedure were used.

Environmental samples from Mediterranean cold seeps with marine AOM associated archaea mainly contained GDGTs with 0 to 2 cyclopentane rings (Pancost *et al.*, 2001). In a study on distinct compartments of AOM-driven carbonate reefs growing in the northwestern Black Sea, GDGTs could only be found in samples when ANME-1 archaea were present, but not when only ANME-2 archaea were found, which led to the conclusion that ANME-2 archaea are not capable of synthesizing internally cyclized GDGT (Blumenberg *et al.*, 2004). Later on in a study on methanotrophic consortia at cold methane seeps, samples associated with ANME-2c were shown to contain relatively high amounts of GDGTs (Elvert *et al.*, 2005). In general, GDGTs are dominant in ANME-1 communities, while in marine ANME-2 and ANME-3 communities archaeol derivatives are most abundant (Niemann and Elvert, 2008; Rossel *et al.*, 2008). Members of the related methanogen order *Methanomicrobiales* produce relatively high amounts of GDGT-0 (Koga *et al.*, 1998; Schouten *et al.*, 2012), whereas *Methanosarcinales* produce no or only minor amounts of GDGTs, mainly GDGT-0 (De Rosa and Gambacorta, 1988; Nichols and Franzmann, 1992; Schouten *et al.*, 2012). Hydroxylated GDGTs seem to be relatively rare. In marine sediment samples the hydroxy-GDGT to total core GDGT ratio has been shown to vary between 1 and 8 % and the dihydroxy-GDGT to total core GDGT ratio is below 2% (Liu *et al.*, 2012). Hydroxylated GDGTs have so far only been identified in the methanogenic Euryarchaeon *Methanothermococcus thermolithotrophicus* (Liu *et al.*, 2012) and in several Thaumarchaeota (Schouten *et al.*, 2012; Sinninghe Damsté *et al.*, 2012). Until now only hydroxylated GDGTs with 0 to 2 cyclopentane rings have been found (Liu *et al.*, 2012; Schouten *et al.*, 2012; Sinninghe Damsté *et al.*, 2012).

Comparing the results obtained in this study and lipid characterizations of marine ANMEs, it is apparent that the ratio of archaeol and GDGTs are distinctive in the different ANME groups: ANME-1 and partially ANME-2c contain substantial amounts of GDGTs and especially in ANME-1, GDGTs are the predominant membrane lipids (Niemann and Elvert, 2008). In contrast to ANME-1, but similar to other ANME-2 and ANME-3, we found that the dominating lipids in the membrane of clade ANME-2d archaea were archaeol variants and not GDGTs. However, about 30% of the membrane lipids in ANME-2d archaea were GDGTs. Most strikingly, the majority of those GDGTs were hydroxylated, which is quite rare and has not been observed for other ANMEs so far.

### Intact polar lipids of *Ca*. Methanoperedens sp

Although intact polar lipids (IPLs) degrade more quickly than core lipids, IPLs are of higher taxonomic specificity and therefore useful to study especially present environments (Ruetters *et al.*, 2002; Sturt *et al.*, 2004). To identify IPLs of *Ca*. Methanoperedens archaea, UHPLC-ESI-MS was performed.

The three most abundant archaeal IPLs detected were archaeol with a dihexose headgroup and hydroxyarchaeol with either a monomethyl phosphatidyl ethanolamine (MMPE) or a phosphatidyl hexose (PH) headgroup. Further headgroups attached to archaeol were monohexose, MMPE, dimethyl phosphatidyl ethanolamine (DMPE), phosphatidyl ethanolamine (PE) and PH. Next to MMPE and PH, hydroxyarchaeol based IPLs also contained dihexose, monopentose, DMPE, PE, pentose-MMPE, hexose-MMPE and pentose-PE. Headgroups of GDGTs were found to be diphosphatidyl glycerol and dihexose phosphatidyl glycerol. The identification of a pentose as a headgroup of hydroyxarchaeol (mass loss of m/z 132) was unexpected. To our knowledge, this is the first description of a pentose as headgroup for microbial IPLs.

ANME-1 archaea mainly produce diglycosidic GDGTs, whereas lipids of marine ANME-2 and ANME-3 are dominated by phosphate-based polar derivatives of archaeol and hydroxyarchaeol (ANME-2: phospatidyl glycerol, phosphatidyl ethanolamine, phosphatidyl inositol, phosphatidyl serine, dihexose; ANME-3: phospatidyl glycerol, phosphatidyl inositol, phosphatidyl serine) (Rossel *et al.*, 2008). Furthermore, marine ANME-2 archaea produce only minor amounts of GDGT-based IPLs and ANME-3 archaea produce no GDGT-based IPLs at all (Rossel *et al.*, 2008). IPLs of ANME-2d archaea can be distinguished from those of ANME-1 archaea by the prevalence of phosphate containing headgroups as well as archaeol and hydroxyarchaeol based IPLs. Furthermore, ANME-2d can be distinguished from other ANME-2 and ANME-3 archaea by the high abundance of dihexose as headgroup, the rare MMPE and DMPE headgroups and putatively also the pentose headgroup, which so far has not been described in the literature. In contrast to ANME-3 archaea, ANME-2d and marine ANME-2 archaea produce GDGT-based IPLs, albeit only in minor amounts.

In marine environments, a variety of archaeal lipids including those identified in ANME archaea can be found, e.g. those of the abundant Thaumarchaeota (GDGTs with hexose or phosphohexose headgroups, Sinninghe Damsté *et al.*, 2012) and uncharacterized archaea (mainly GDGTs with glycosidic headgroups and in subsurface sediments also archaeol with glycosidic headgroups, Sturt *et al.*, 2004; Lipp *et al.*, 2008). In freshwater environments, IPLs of methanotrophic archaea have hardly been studied. Two studies on peat samples identified GDGTs with a glucose or glucuronosyl headgroup (Liu et al., 2010) and with a hexose-glycuronic acid, phosphohexose, or hexose-phosphoglycerol head group (Peterse et al., 2011). GDGTs with a hexose-phosphoglycerol head group were also identified in our study for ANME-2d archaea. Therefore, ANME-2d together with other archaea might be part of the peat microbial community based on the IPL profile. Using DNA biomarkers, most notably the 16S rRNA gene, *Ca*. Methanoperedens sp. has been detected in various peat ecosystems (Cadillo-Quiroz *et al.*, 2008; Zhang *et al.*, 2008; Wang *et al.*, 2019).

The related order *Methanosarcinales* mainly produce archaeol and hydroxyarchaeol with the headgroups glucose, phosphatidyl glycerol (only *Methanosarcinaceae*), phosphatidyl inositol, phosphatidyl ethanolamine, galactose (only *Methanosaetaceae*) (Koga *et al.*, 1998). On the other hand, members of the related order *Methanomicrobiales* contain GDGT-0 and archaeol with the lipid headgroups glucose, galactose, phosphatidyl aminopentanetetrols, phosphatidyl glycerol (Koga et al., 1998). Therefore, IPLs from *Ca*. Methanoperedens sp. differ from methanogen IPLs by the high abundance of dihexose, MMPE and phosphatidyl hexose as lipid headgroup and the absence of the quite common headgroup phosphatidyl serine.

### Incorporation of carbon derived from methane and bicarbonate in lipids

We were not only interested in characterizing the lipids of *Ca*. Methanoperedens sp., but also in answering the question if the organism incorporates carbon derived from methane or from dissolved inorganic carbon (DIC) in its lipids. In a labelling experiment from 2006 with an ANME-2d enrichment culture, incorporation of carbon derived from methane could hardly be detected for archaeal lipids (Raghoebarsing *et al.*, 2006). To establish the carbon sources for *Ca*. Methanoperedens sp. we incubated the enrichment culture with ^13^C labelled bicarbonate and methane and analysed lipid extracts for δ^13^C depletion by GC-IRMS (Fig. 2).

**Figure 2:**
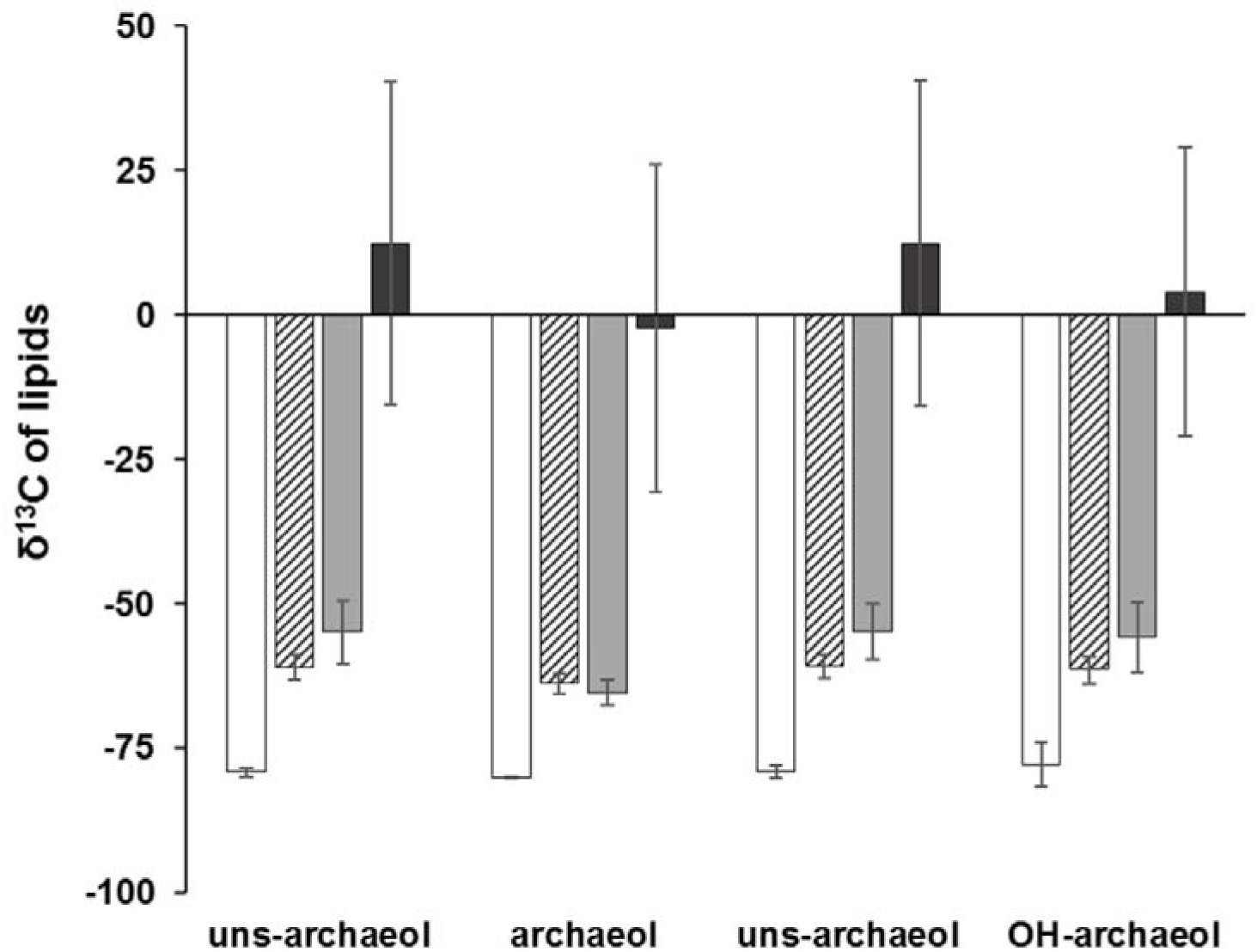
δ^13^C values of *Ca*.Methanoperedens sp. lipids after batch cultivation with labelled bicarbonate or methane. ANME-2d reactor material originating from the Ooijpolder was incubated in anaerobic batch cultures with either 13C labelled bicarbonate for three days (striped columns) or ^13^C labelled methane for one (light grey columns) or three days (dark grey columns) and analysed via GC-IRMS. Controls contained only non-labelled carbon sources (white columns). Incubations were performed in triplicates, error bars = standard deviation. Peak identification was conducted with the help of GC-MS analysis of the same samples, showing that lipid extracts contained archaeol, hydroxyarchaeol and two monounsaturated archaeols (Fig. 1). Uns-archaeol = monounsaturated archaeol.

Analysis of the isotopic composition of archaeol and its derivatives showed that ANME-2d archaea incorporated carbon derived from both methane and bicarbonate into their lipids. However, the main carbon source for biomass production seemed to be methane and not DIC as the former shows more label in the archaeal lipids. However, it has to be considered that the cultures to which ^13^C labelled bicarbonate was added did not exclusively contain ^13^C-DIC. About half of the DIC in the cultures derived from ^12^C-CO_2_ dissolved in the medium after gassing with a mixture of 10% CO_2_/90% Argon gas (calculations in the methods part). Considering this, the δ^13^C values of the archaeol isomers without ^12^C-DIC in the incubations would vary most probably between −40 and −60‰. Nevertheless, the respective lipids were still quite depleted in δ^13^C in comparison to the incubations with labelled methane (−2 to 12‰; 3 days incubation). Therefore, we concluded that mainly methane and not DIC is incorporated in the lipids of *Ca*. Methanoperedens sp.. Supporting this result, cultures containing marine ANME-1 and ANME-2 were shown to incorporate carbon derived from labelled methane into archaeol, monounsaturated archaeol and biphytanes (Blumenberg *et al.*, 2005). In another study it was found that ANME-1 archaea assimilated primarily inorganic carbon (Kellermann *et al.*, 2012). Incubations with sediments containing ANME-1, 2a & 2b archaea showed that both, labelled methane and inorganic carbon, were incorporated into the archaeal lipids (Wegener *et al.*, 2008). Incubations with freshwater sediments including ANME-2d archaea followed by RNA stable isotope probing demonstrated that those microbes mainly incorporated methane into their lipids but may have the capability of mixed assimilation of CH_4_ and dissolved inorganic carbon (Weber *et al.*, 2017). Our data confirmed that ANME-2d archaea are capable of mixed assimilation of CH_4_ and DIC, but that methane is the preferred carbon source.

## Conclusion

In this study, we analysed the lipids from the main player in nitrate AOM, *Ca.* Methanoperedens sp. We found several lipid characteristics that enable distinction between ANME-2d and other ANME groups (Table 2).

**Table 2:** Lipids of different ANME groups. For ANME-2d lipid analysis we used *Ca*. Methanoperedens sp. enriched bioreactor material. For the other ANME groups information was based on publication about the specific lipid characteristic (Blumenberg *et al.*, 2004; Niemann and Elvert, 2008) or ^13^C labelling experiments (Blumenberg *et al.*, 2005; Wegener *et al.*, 2008; Kellermann *et al.*, 2012). GDGT: glycerol dialkyl glycerol tetraether, PE: phosphatidyl ethanolamine, MMPE: monomethyl phosphatidyl ethanolamine, DMPE: dimethyl phosphatidyl ethanolamine, PG: phosphatidyl glycerol, MH: monohexose, DH: dihexose, PH: phosphatidyl hexose, PC: phosphatidyl choline.

|  | ANME-1 | ANME-2a/b | ANME-2c | **ANME-2d** | ANME-3 |
| --- | --- | --- | --- | --- | --- |
| Environment | marine | marine | marine | **freshwater** | marine |
| core lipids | GDGT | (OH-) archaeol | (OH-) archaeol, GDGTs | **(OH-) archaeol, (OH)-GDGTs** | (OH-) archaeol |
| Sn-2-OH-archaeol / archaeol ratio | 0 - 0.8 | 1.1 - 5.5 | 1.1 - 5.5 | **0.1 - 0.3** | 1.1 - 5.5 |
| IPLs | GDGT + dihexose | (OH-) Archaeol + PG, PE, PH, PS, dihexose | (OH-) Archaeol + PG, PE, PH, PS, Dihexose | **(OH-) archaeol + dihexose, hexose, pentose, PH, PE, MMPE, DMPE** | (OH-) Archaeol + PG, PH, PS |
| Main carbon source | DIC |  |  | **CH<sub>4</sub>** |  |

ANME-2d archaea therefore can be distinguished from ANME-1 by the higher ratio of archaeol and hydroxyarchaeol instead of GDGTs as well as phosphate containing headgroups. Furthermore, ANME-2d can be distinguished from other ANME-2 and ANME-3 archaea by the high abundance of dihexose as headgroup, the rare MMPE and DMPE headgroups and putatively also the pentose headgroup, which so far has not been described in the literature. The appearance of a monopentose as headgroup of ANME-2d lipids is an interesting observation and might be further analysed in the future. In contrast to other ANME groups ANME-2d archaea have been shown to produce relatively rare hydroxylated GDGTs.

ANME groups do not only differ in their membrane lipids itself, but also in the way they incorporate carbon into their biomass. For ANME-1 it has been shown that primarily carbon derived from DIC is incorporated into the lipids (Kellermann *et al.*, 2012). In case of ANME-2d archaea, we were able to demonstrate that both, carbon derived from DIC and from methane, are incorporated into their lipids, with methane as the preferred carbon source.

## Acknowledgements

CUW and MSMJ were supported by the Nederlandse Organisatie voor Wetenschappelijk Onderzoek through the Soehngen Institute of Anaerobic Microbiology Gravitation Grant 024.002.002 and the Netherlands Earth System Science Center Gravitation Grant 024.002.001. MSMJ was supported by the European Research Council Advanced Grant Ecology of Anaerobic Methane Oxidizing Microbes 339880. JK was supported by the Netherlands Earth System Science Center Gravitation Grant 024.002.001 and the Deutsche Forschungs Gesellschaft (DFG) Grant KU 3768/1-1. SB and CW were supported by the Nederlandse Organisatie voor Wetenschappelijk Onderzoek through Grant ALWOP.293. We thank Michel Koenen for IPL analysis, Ronald van Bommel for technical assistance with the GC-IRMS, Denise Dorhout and Monique Verweij for technical assistance with GC-MS and UHPLC systems. Moreover, we thank Annika Vaksmaa for supplying bioreactor material, Theo van Alen for technical assistance with metagenomics sequencing and Jeroen Frank for helping with metagenomics analysis.

